# *PSGfinder*: fast identification of genes under divergent positive selection using the dynamic windows method

**DOI:** 10.1101/193722

**Authors:** Joël Tuberosa, Juan I. Montoya-Burgos

## Abstract

**Summary:** Orthologous genes evolving under divergent positive selection are those involved in divergent adaptive trajectories between related species. Current methods to identify such genes are complex and conservative or present some imperfections, limiting genome-wide searches. We present a simple method, Dynamic Windows, to detect regions of protein-coding genes evolving under divergent positive selection. This method is implemented in *PSGfinder*, a user-friendly and flexible software, allowing rapid genome-wide screenings of regions with a dN/dS >1. *PSGfinder* additionally includes an alignment cleaning procedure and an adapted multiple comparison correction to identify significant signals of positive selection.

**Availability and Implementation:** *PSGfinder* is a software that implements the DWin method, is written in Python and is freely available with its documentation at: <u>https://genev.unige.ch/research/laboratory/Juan-Montoya</u> or at: <u>https://github.com/joel-tuberosa/psgfinder</u>

## 1 Introduction

Positive selection is the process in which alleles that have a positive effect on the fitness of the individuals bearing them will spread into the population and reach fixation faster than alleles with no such effect. Codon-based likelihood models have been widely used to detect positive selection in protein-coding genes, among which the branch-site test (Zhang, et al., 2005). This test generally requires more than four orthologous sequences; its power and computation time increases with the number of sequences. The branch-site test is often considered to be too conservative (Zhang, et al., 2005) with a significance threshold exaggeratedly stringent (Nickel, et al., 2008). The Variation Clusters is an alternative fast method but applies only to moderately diverged sequences (Wagner, 2007) and suffers from the observation that the signatures of positive and purifying selection can be confounded (McFerrin and Stone, 2011).

Divergent positive selection refers to the process in which the orthologous genes of two closely related species undergo positive selection along divergent adaptive trajectories. This typically occurs when sister species adapt to different environments. When comparing the orthologous protein-coding genes of two closely related species, regions evolving under divergent positive selection are classically characterized by a higher non-synonymous mutation rate (dN) than the synonymous mutation rate (dS), that is, with a dN/dS ratio >1. The sliding window approach can rapidly scan gene regions of pairwise alignments to identify windows showing a dN/dS >1 indicative of positive selection (e. g. Montoya-Burgos, 2011). However, this approach has limitations: no general criterion to define window size and window shift; when many windows are analyzed per gene, false positives become difficult to avoid; windows can have dS=0 excluding dN/dS calculation; the window dS accuracy decreases with decreasing window size (Montoya-Burgos, 2011; Schmid and Yang, 2008).

Here, we present a new method to detect divergent positive selection, Dynamic Windows (DWin), that solves the above-mentioned limitations, and the software that implements it, *PSGfinder.*

## 2 Methods and Implementation

DWin is based on the observation that positively selected sites are generally clustered along the gene as they belong to a same functional motif of the encoded protein. Consequently, in pairwise alignment of orthologous genes, regions (windows) under divergent positive selection will show a high dN. To have a reasonable chance to pass the significance threshold, we inferred that windows must be at least 3 codons long (criterion 1) and contain at least 3 amino acid differences (criterion 2). Thus, for each pairwise alignment, DWin will consider all the windows that start and end with an amino acid difference and that fulfill the two criteria. Windows are allowed to overlap partially or completely. To avoid low dS accuracy in small windows or dS=0, we introduce the use of the entire gene’s dS but the window’s dN to calculate the dN/dS per window, with a dN/dS significantly >1 indicating positive selection.

*PSGfinder* implements DWin and performs two additional operations: alignment cleaning and a correction for multiple comparisons. We developed four filters for detecting misaligned regions as they can generate false signals of positive selection. First, regions with numerous interspersed gaps often result from the alignment of non-homologous regions, so only regions with >10 codons without gaps are kept; otherwise they are masked. Second, misaligned regions generally show excessive dS, so alignment regions flanked by gaps (or masks) that display a dS >1 are masked (excessively high for closely related species). Third, misaligned regions also display an excessive concentration (a cluster) of non-synonymous substitutions. To detect such clusters, the alignment is translated into amino acids (a.a.) and contiguous (≥2) variable a.a. positions are defined as cluster sub-units; when sub-units are separated by <4 positions they form a cluster. Clusters containing >8 variable a.a. positions are masked. Fourth, gene pairs displaying an overall dS >1 are considered as non-homologs and are discarded. All these threshold values are user-modifiable.

*PSGfinder* uses the Fisher exact test to assess if dN/dS is significantly >1. To overcome false positives when generating more and more windows and to account for windows overlap, we adapted a Bonferroni correction (to the p-value threshold) in which the number of non-overlapping tests per gene is determined as follows: Each pairwise alignment is translated into a.a. For each group (*k*) of overlapping windows encompassing *Sk* sites with an a.a. mismatch and containing *Nk* windows with each *Sw* sites with an a.a. mismatch, the number of non-overlapping tests (*Tk*) is approximated by:

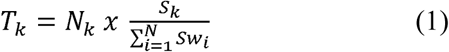

For each gene, the number of non-overlapping tests (*T*) is the sum of all *Tk. T* is used to correct the significance threshold.

*PSGfinder* input data is a collection of FASTA formatted pairwise alignments of the orthologous coding sequences of the genes to be analyzed. An example is provided in Supplementary information. *PSGfinder* is written in Python and is executable in Windows, Linux and MacOS. *PSGfinder* uses one public dependency, PAML (Yang, 2007) and two downloadable Python modules, numpy (<u>http://www.numpy.org/</u>) and fisher (<u>https://pypi.python.org/pypi/fisher/</u>)

## 3 Example usage

*PSGfinder* has been implemented in the comparative transcriptomics work by Weber, et al. (2017). To uncover genes involved in reproductive barriers between species, these authors recovered 12,229 orthologous protein-coding genes between two brittle star species (*Ophioderma*). By using *PSGfinder,* they identified 48 genes that displayed regions with divergent positive selection, from which two genes were of interest as they are involved in reproduction. These genes were confirmed to evolve under positive selection when analyzed more deeply with additional data and using the classical branch-site model approach. As a detailed example on how to use *PSGfinder* we provide the data input of Weber *et al.* (2017) and the command instructions in the documentation of *PSGfinder*.

## 4 Significance and conclusions

The DWin method to detect gene regions undergoing divergent positive selection requires only the coding sequence of the orthologous genes of the two focus species and uses the standard dN/dS ratio statistics to evaluate selection. DWin examines only promising candidate windows, minimizing the correction for multiple comparisons and generating results rapidly. *PSGfinder* implements DWin in a user friendly, flexible and computationally fast manner allowing the analysis of genomic sets of protein-coding genes. We emphasize that our method is complementary to existing approaches and is useful to perform fast genome-wide screens to identify candidate genes that can be analyzed further with classical approaches.

## Acknowledgments

We acknowledge Dr. L. Capello and Dr. J.-L. Falcone for useful advises. We thank Dr. A. Weber for providing data in the example usage.

